# Live Zika virus chimeric vaccine candidate based on a yellow fever 17-D attenuated backbone

**DOI:** 10.1101/272625

**Authors:** Franck Touret, Magali Gilles, Raphaelle Klitting, Fabien Aubry, Xavier de-Lamballerie, Antoine Nougairède

**Affiliations:** Unité des Virus Émergents (UVE: Aix-Marseille Univ – IRD 190 – Inserm 1207 – IHU Méditerranée Infection), Marseille, France

## Abstract

Zika virus (ZIKV) recently dispersed throughout the tropics and sub-tropics causing epidemics associated with congenital disease and neurological complications. There is currently no commercial vaccine for ZIKV. Here we describe the initial development of a chimeric virus containing the prM/E proteins of a ZIKV epidemic strain incorporated into a yellow fever 17-D attenuated backbone. Using the versatile and rapid ISA (Infectious Subgenomic Amplicons) reverse genetics method, we compared different constructs and confirmed the need to modify the cleavage site between the pre-peptide and prM protein. Genotypic characterization of the chimeras indicates that emergence of compensatory mutations in the E protein is required to restore virus replicative fitness. Using an immunocompromised mouse model, we demonstrate that mice infected with the chimeric virus produced levels of neutralizing antibodies close to those observed following infection with ZIKV and that pre-immunized mice were protected against viscerotropic and neuroinvasive virus following challenge with a heterologous strain of ZIKV. These data provide a sound basis for the future development of this ZIKV vaccine candidate.

## Introduction

Zika virus (ZIKV; family *Flaviviridae*, genus *Flavivirus)* is a single-stranded positive-sense enveloped RNA virus. Its genome of 10.8 kb encodes a single polyprotein which is processed into three structural proteins (C, PrM and E), and seven nonstructural proteins (NS1, NS2A, NS2B, NS3, NS4A, NS4B and NS5) by viral and host proteases^1^. Phylogenetic studies showed that all ZIKV strains characterized so far belonged among two distinct lineages (African and Asian) based on the initial geographic distribution of this virus^2^. ZIKV is a mosquito-borne flavivirus transmitted primarily by *Aedes spp.* mosquitoes^3^.

Long considered to cause mild disease in humans, this arbovirus remained relatively unstudied until 2007, when it provoked a large outbreak in Micronesia^4^. Subsequently, several outbreaks occurred in different Pacific Ocean islands including French Polynesia in 2013, where it was associated with an increase of Guillain-Barré syndrome^5^. ZIKV then spread to the American continent causing major outbreaks in Central/South America and the Caribbean and was linked with an increase of congenital neurological complications. Sexual transmission of ZIKV was also reported^6^. There is currently no commercial antiviral drug or vaccine for this virus^7^.

Several approaches are now available with which to develop inactivated^8^ and recombinant (DNA-^9^ or RNA-based^10^) ZIKV vaccines. However, live attenuated vaccines have several advantages including reduced costs and single dose induction of long-term immunity^11^. Several groups developed live ZIKV vaccine candidates by deletions in the 3’ untranslated region of the viral genome^12,13^. More recently, a chimeric ZIKV vaccine candidate based on the Japanese encephalitis virus live-attenuated train SA14-14-2 was reported^14^. The chimeric approach had been used since the late 1990s to develop vaccine candidates against several health-threatening flaviviruses including West-Nile virus, Japanese encephalitis virus and all serotypes of the dengue virus^15–17^. This approach consists of incorporating prM/E of a pathogenic flavivirus in a backbone of a licensed live-attenuated vaccine strain. Indeed, E protein is prominently exposed at the surface of viral particles and is *de facto* the major determinant of viral antigenicity^1^. In almost all cases, the well-characterized live attenuated 17-D strain used to prevent yellow fever virus (YFV) infections has been used as the genetic backbone. Some of these live-attenuated vaccines are currently commercially available^18,19^.

Here we describe the development of a chimeric virus harboring the prM/E of an epidemic ZIKV (H/PF/2013) strain and the 17-D vaccine strain as the genetic backbone. The user-friendly and rapid ISA (Infectious Subgenomic Amplicons) reverse genetics method was used to generate this chimeric virus^20^. Finally, *in cellulo* and *in vivo* characterization of this strain demonstrated its potential to become a live-attenuated vaccine candidate.

## Results

### Design and rescue of chimeric viruses

The chimeric viruses were constructed using the yellow fever 17-D vaccine strain as a genetic backbone and by replacing the prM/E of this vaccine strain by those of the Asian ZIKV PF epidemic strains. Three different constructs, designated A, B and C, were constructed using variable sites flanking ZIKV prM/E coding sequences (Fig. 1). Construct A harbored the prepeptide and cleavage site before prM from the 17-D vaccine strain. Construct B harbored the pre-peptide from the 17-D vaccine strain and cleavage site before prM from ZIKV. Construct C harbored the pre-peptide and cleavage site before the prM from ZIKV. They all contained the cleavage site of the 17-D vaccine strain between E and NS1 proteins.

**Figure 1.**
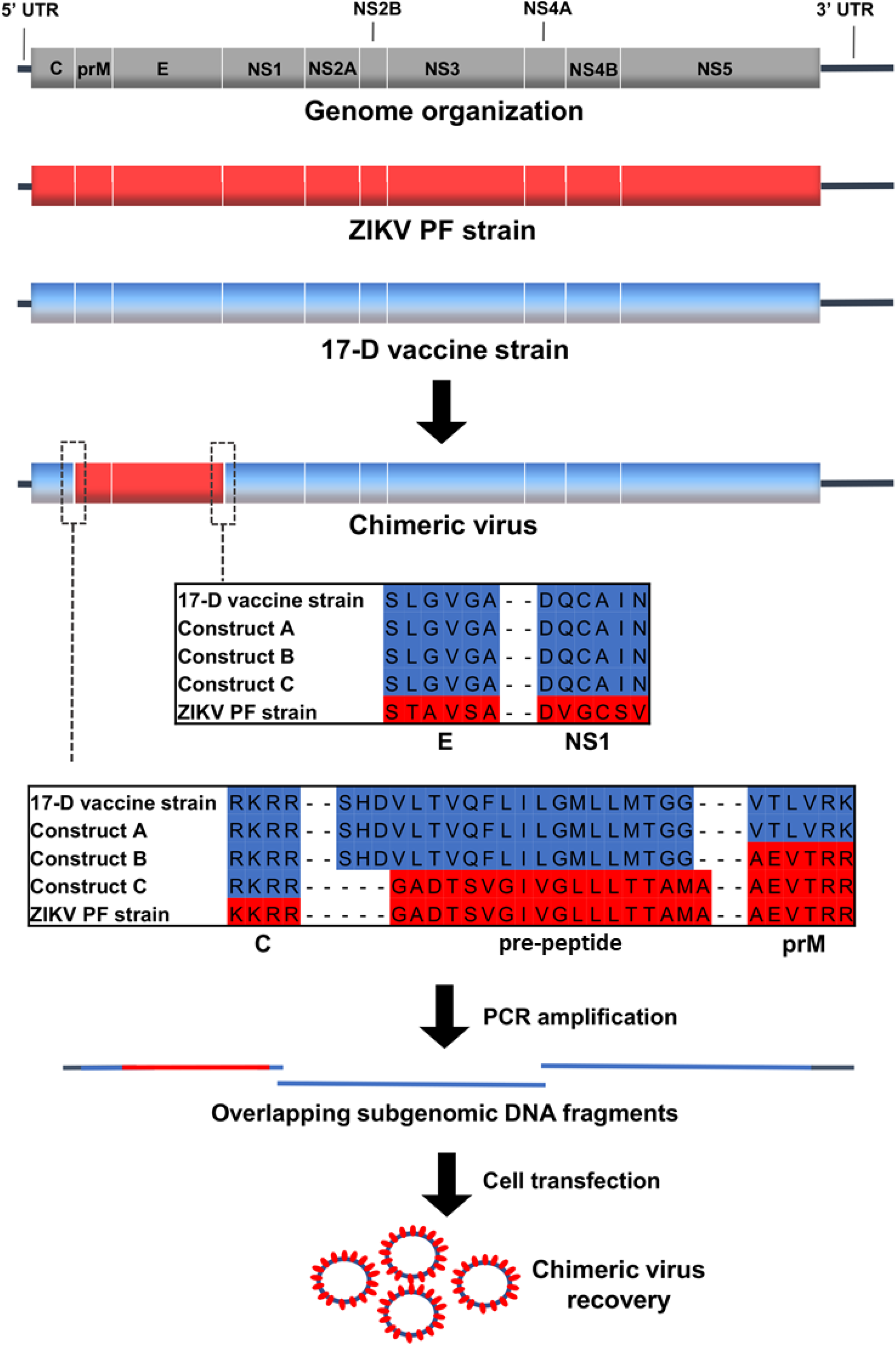
Schematic representation of design and recovery strategies used to generate chimeric viruses. We recovered infectious virus with construct C only. The two cleavage sites are enlarge in boxes with the amino acid alignment represented with separation between different proteins.

The ISA procedure was used to rescue the viruses: Three overlapping amplicons that contain the complete genome flanked at its 5’ and 3’ extremities respectively by the human cytomegalovirus promoter (pCMV) and the hepatitis delta ribozyme followed by the simian virus 40 polyadenylation signal (HDR/SV40pA) were transfected into a mix of HEK-293/BHK21 cells. Because the first amplicon contains the entire structural coding region, it was only necessary to exchange the first amplicon of our already functional yellow fever 17-D vaccine strain reverse genetic system to attempt replicative virus production.

For each construct, we performed two independent cell transfection experiments of five replicates. After an incubation of six days, cell supernatant media were passaged four times in Vero-E6 cells. Virus replication was assessed in cell supernatant medium at the last passage (called passage #4) using a real-time quantitative RT-PCR assay. No viral replication was detected for constructs A and B. In contrast, we detected, with construct C, virus replication in one well (1/5) in both independent transfection experiments. These results highlighted the fact that the choice of the nature of the pre-peptide and cleavage site between the capsid and prM proteins is a crucial parameter when designing chimeric flaviviruses. During the first cell transfection experiment, we obtained at passage #4 high amounts of viral genome copies (1.78 e+9 viral RNA copies/mL). This virus was designated CH-17-D/ZIKV and used for *in cellulo* and *in vivo* characterizations. Surprisingly, during the second experiment, we detected very low quantities of viral genome at passage #4 (3.57 e+3 viral RNA copies/mL). This virus was designated CH-17-D/ZIKV*. We then performed four additional passages using the same procedure. Quantities of viral genomes in cell supernatants media were assessed from passage #1 to #8 and compared with that of CH-17-D/ZIKV (Fig. 2). We found that amounts of viral genomes for CH-17-D/ZIKV reached a plateau at passage #2. In contrast, we observed an increase in the production of CH-17-D/ZIKV* from passage #6 to passage #8 where amounts of viral genome reached values similar to those observed with CH-17-D/ZIKV (2,67e+9 viral RNA copies/mL) (Fig. 2).

**Figure 2.**
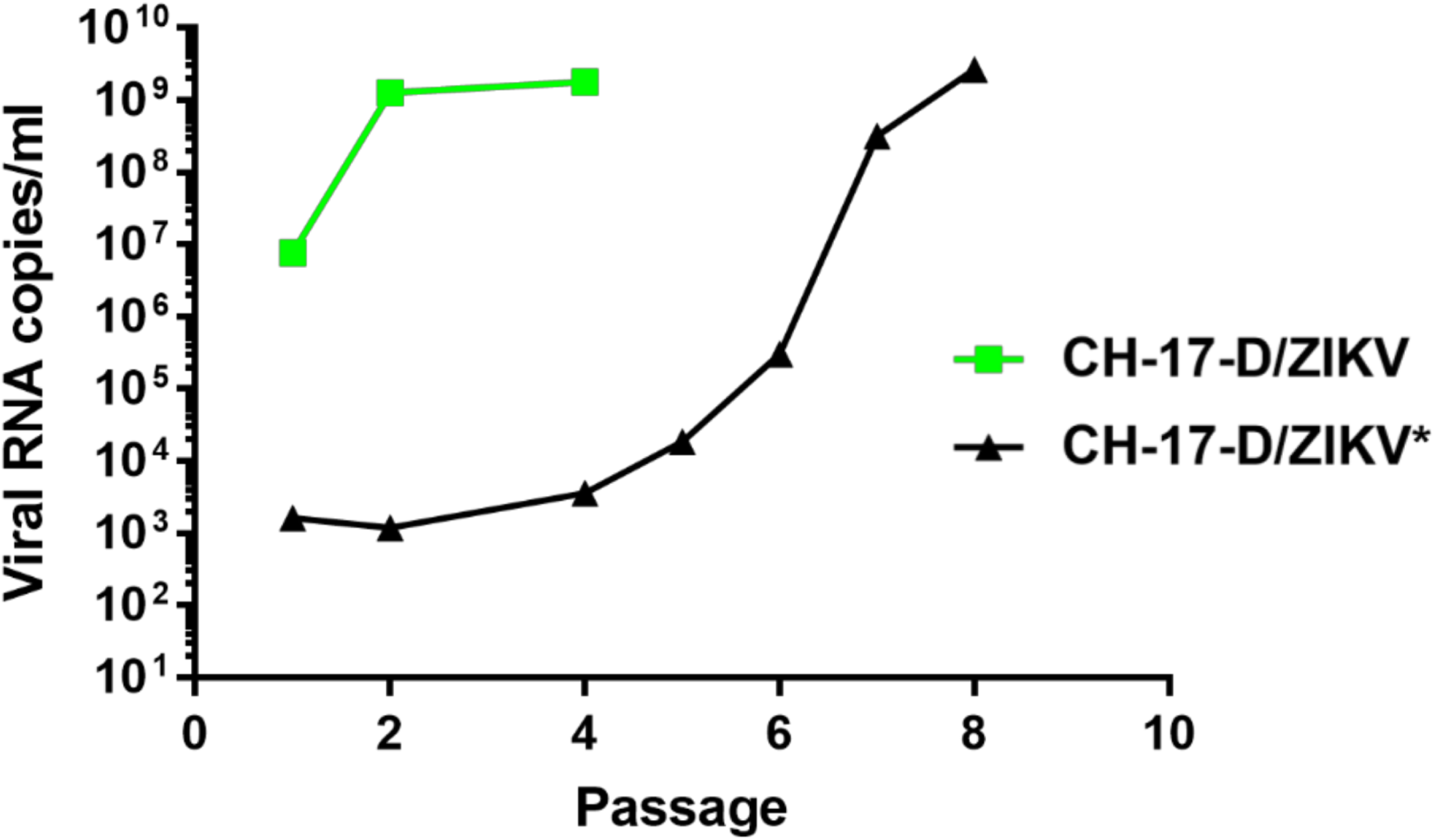
Evolution of viral production of chimeric viruses during serial passage that followed cell transfection. A mix of BHK-21/HEK-293 cells was transfected. Cell supernatant media were then passaged 4-8 times in Vero-E6 cells. Viral production in cell supernatant medium was assessed using a real time quantitative RT-PCR assay.

### CH-17-D/ZIKV genotypic characterization

In order to find the genotypic determinants associated with the difference of viral replication observed between CH-17-D/ZIKV and CH-17-D/ZIKV*, the complete genome of CH-17-D/ZIKV was obtained at passage #2 and #4 and compared with the sequence of the original construct. Only five substitutions were detected at passage #2 of which two were non-synonymous confirming the genome integrity of this strain (Table 1). Four of them were already fixed or almost fixed. At passage #4, all these mutations were fixed and no additional mutations were found. Interestingly, both non-synonymous mutations are located in domain II of E protein, respectively at residues E255 and E285^21^. Consequently, we determined the sequence of the 5’ region of the viral genome of CH-17-D/ZIKV* (until NS1 coding region) at passage #4 and the complete genome sequence of CH-17-D/ZIKV* at passage #8 (Table 1). While only one transitory substitution was detected at passage #4, all the mutations found with CH-17-D/ZIKV were detected at passage #8 including the two non-synonymous mutations located in E coding region. This high level of parallel evolution associated with the chronology of events strongly suggests that these five mutations are associated with the increase of replicative fitness observed with both viruses.

**Table 1:** Mutations detected during the passages that followed cell transfection of chimeric viruses. Only consensus mutations (frequency >50%) are shown. n.a: not available; n.d: not detected

| Chimeric virus | Nucleotide position | Frequency at #P2 | Frequency at #P4 | Frequency at #P8 | Frequency at #P10 | Region | Nucleotide change | aa change |
| --- | --- | --- | --- | --- | --- | --- | --- | --- |
| CH-17-D/ZIKV | 291 | 100% | 100% | 100% | 100% | C | A>G | - |
|  | 1625 | 66% | 90% | 92% | 94% | E | T>C | V>A |
|  | 1706 | 100% | 100% | 100% | 100% | E | G>T | G>V |
|  | 2514 | 100% | 100% | 100% | 79% | NS1 | A>G | - |
|  | 4482 | 100% | 100% | 100% | 100% | NS2B | A>G | - |
| CH-17-D/ZIKV* | 291 | n.a | n.d | 70% | n.a | C | A>G | - |
|  | 1303 | n.a | 100% | n.d | n.a | E | C>T | H>Y |
|  | 1625 | n.a | n.d | 68% | n.a | E | T>C | V>A |
|  | 1706 | n.a | n.d | 69% | n.a | E | G>T | G>V |
|  | 2514 | n.a | n.d | 69% | n.a | NS1 | A>G | - |
|  | 4482 | n.a | n.a | 56% | n.a | NS2B | A>G |  |

### CH-17-D/ZIKV initial characterization

To confirm the presence of the ZIKV E protein in Vero-E6 cells infected by the chimeric virus, we performed an indirect immunofluorescence assay using a specific ZIKV immune serum as the primary antibody (Fig. 3A). ZIKV PF and the 17-D vaccine strains were used as positive and negative controls. As expected, no fluorescence was observed with the 17-D vaccine strain and positive cells were observed at day 2 and 5 post-infection with both chimeric and ZIKV strains confirming that the ZIKV E protein is expressed in infected cells. At day 2 postinfection, the number of positive cells with ZIKV is higher than with CH-17-D/ZIKV in agreement with growth replication kinetics in VeroE6 cells. Since a cytopathic effect was observed with the ZIKV strain at day 5, the number of positive cells was lower with this virus. Viability assays in Vero-E6 cells confirmed this observation: the CH-17-D/ZIKV virus is less cytopathic (mean value: 73% of cell viability) at day 5 post-infection than the ZIKV (mean value: 49% of cell viability) (Supplemental Fig. 1). We then performed comparative growth kinetics of these viruses in three different cell lines (HUH7.5, HEK-293 and Vero-E6). Cell supernatant media were harvested at different time points after infection to assess the amount of viral RNA (Fig. 3B, 3C and 3D). Similar growth kinetics curves were observed for all viruses in HUH7.5. In Vero-E6 cells, higher amounts of viral genome were not found in cell supernatants until day 5 post-infection with the chimeric virus. In HEK-293 cells, the chimeric virus had similar behavior to that of the 17-D vaccine strain.

**Figure 3.**
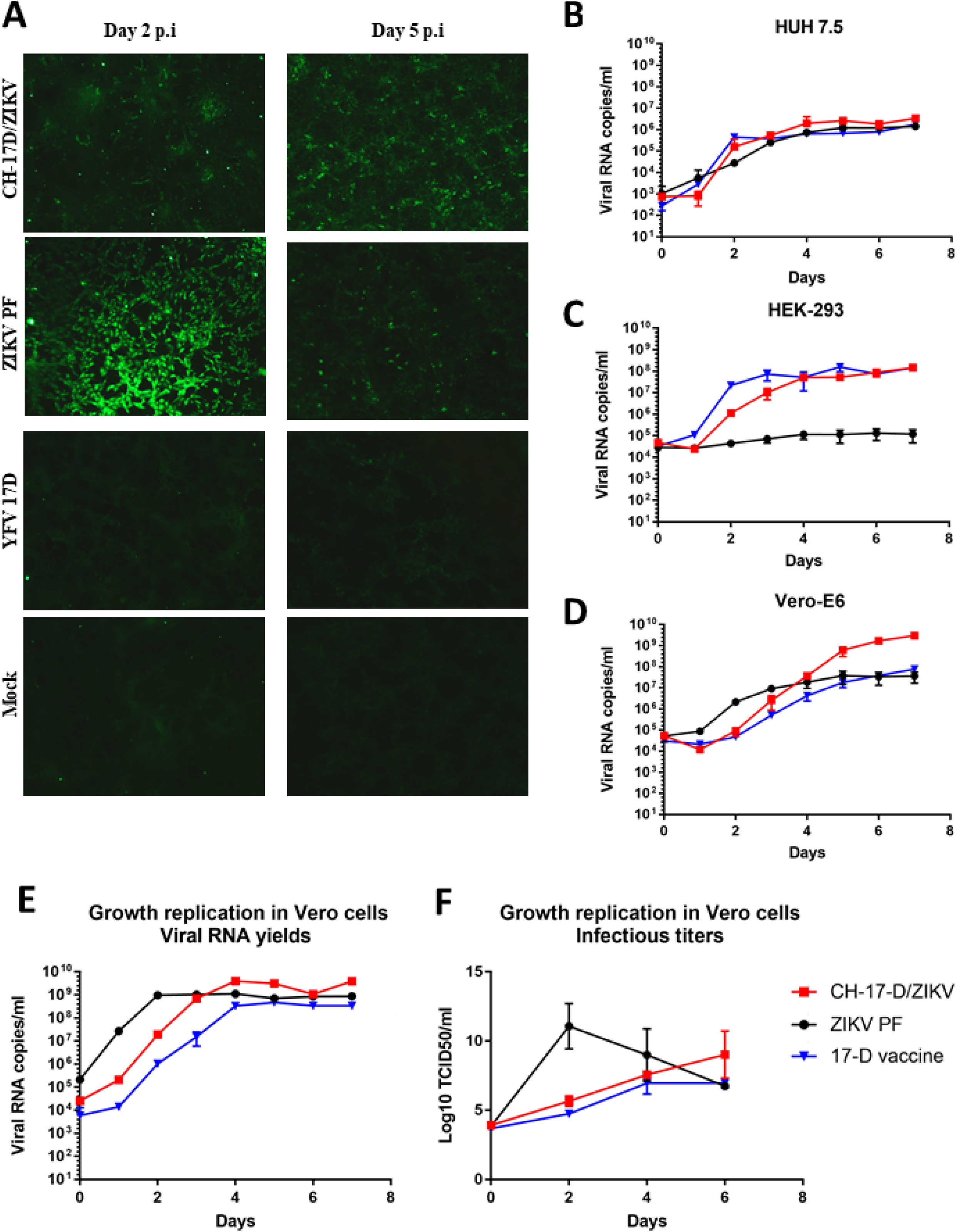
CH-17-D/ZIKV *in cellulo* characterization. **Panel A:** Expression of the ZIKV E protein in Vero-E6 was confirmed at day 2 and 5 postinfection using an indirect immunofluorescence assay using a specific ZIKV immune serum as the primary antibody. Uninfected cells (mock) and cells infected by ZIKV and the 17-D vaccine strain were used as controls. **Panel B/C/D:** Comparative growth kinetics of CH-17-D/ZIKV, ZIKV 17-D vaccine strains HUH 7.5 (**B**), HEK-293 (**C**) and Vero E-6 (**D**). **Panel E/F**: Comparative growth kinetics of CH-17-D/ZIKV, ZIKV 17-D vaccine strains in Vero cells. Cell supernatant media were harvested at different time points after infection to assess the amount of viral RNA using a real time quantitative RT-PCR assay (**panel E**; expressed as mean ±SD) and the infectious titers using a TCID50 assay (**panel F**; expressed as mean ±SD).

### CH-17-D/ZIKV characterization in Vero cells

Since Vero cells are widely used for vaccine production^22^, we characterized CH-17-D/ZIKV in this cell line. Because CH-17-D/ZIKV is already adapted at passage #4 (see above), we used cell supernatant from this passage to perform growth kinetics in Vero cells. Cell supernatant media were harvested at different time points after infection to assess infectious titers (TCID50 assay) and the amount of viral RNA (Fig. 3E/F). The results showed that these cells enabled the production of highly infectious viral particle at day 6 post-infection. We also studied the genetic stability of CH-17-D/ZIKV by performing 6 additional passages in Vero cells and the complete genome sequence was obtained at passage #8 and #10 (Table 1). Our findings revealed remarkable genetic stability since all mutations at passage #4 remained stable and no additional mutation was detected.

### CH-17-D/ZIKV *in vivo* characterization

Because ZIKV and 17-D vaccine strain do not replicate in immunocompetent mice, we used immunocompromised mice as model to study the chimeric virus *in vivo.* Each time animals were immunized or infected, they were transiently immunocompromised following a two-step inoculation of anti-IFNAR antibody ^23–25^ as described in the Methods section.

Six groups of four mice were inoculated with two different dosages of CH-17-D/ZIKV, ZIKV,17-D vaccine strain to assess antibody production (Fig. 4A). A control group (mock) of four mice were inoculated with PBS. Twenty-one days after immunization, mice were sacrificed and their sera were tested for the presence of antibodies to ZIKV and YFV (Fig. 4B and 4C) using a viral RNA Yield Reduction Neutralization Test (YRNT; see Methods). As a result, we demonstrated that immunization with the chimeric virus induced production of neutralizing antibodies against ZIKV confirming the starting hypothesis of this study. We detected a slightly higher level of neutralizing antibodies when mice were infected with ZIKV. In all cases, both dosages used induced comparable neutralizing titers. Consistently with previous studies, mice immunized with the 17-D vaccine strain did not produce antibodies against ZIKV. Indeed, based on the amino acid sequence divergence of antigenic proteins, it is well established that no cross-neutralizing activity exists between these two distant flaviviruses^26^. As expected, mice immunized with the 17-D vaccine strain produced high levels of neutralizing antibodies against YFV while those infected with ZIKV did not produce any antibodies against YFV. Interestingly, immunization with CH-17-D/ZIKV induced production of neutralizing antibodies against YFV. This result demonstrated the immunogenicity of the viral proteins encoded by the 17-D vaccine strain backbone. We also attempted to isolate chimeric virus from animal blood samples to assess the ability of the chimeric virus to replicate *in vivo.* At day 2 and 3 post immunization, to avoid the possibility of isolating residual virus from the immunization, we collected a blood drop from the mice tail and found two positive samples (one at each day) (Supplemental Fig. 2). These findings suggest that CH-17-D/ZIKV is able to replicate in mice since comparable neutralizing titers were measured with all mice immunized.

**Figure 4.**
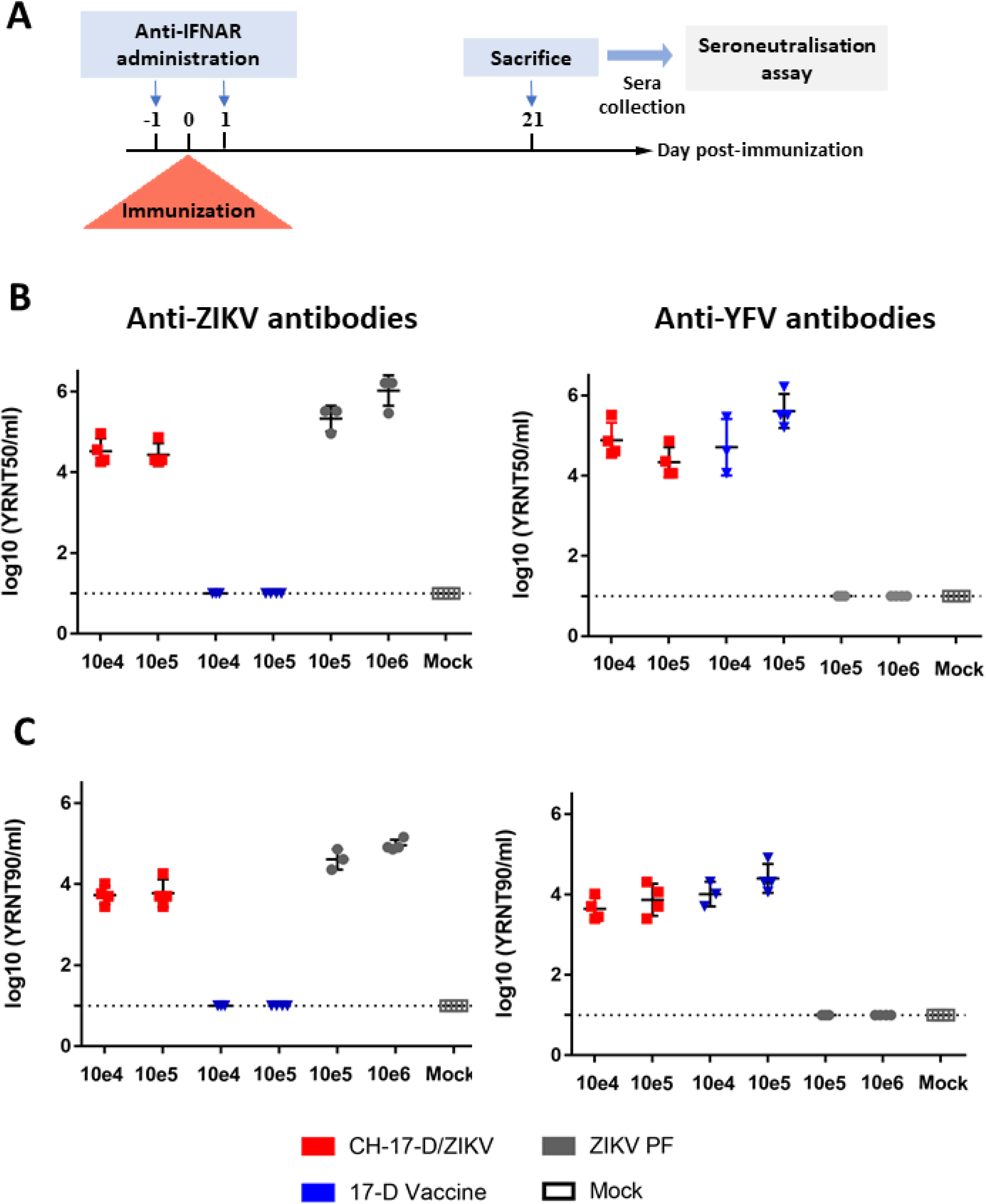
Neutralizing antibody titers of transitory immunocompromised mice at day 21 post-immunization. **Panel A**: Experimental timeline. **Panel B/C**: Groups of four mice were immunized with two doses of CH-17-D/ZIKV, ZIKV and the 17-D vaccine strain (from 10e4 to 10e6 TCID50). Twenty-one days later, sera from mice were tested for the presence of antibodies to ZIKV and YFV using a viral RNA Yield Reduction Neutralization Test. Results are expressed as individual log of YRNT50 (Panel B) and YRNT90 titers (**Panel C**) with mean values ±SD represented respectively by black lines and error-bars.

In another experiment, we assessed protection against subsequent infection by wild-type ZIKV following immunization with CH-17-D/ZIKV or 17-D vaccine strain (Fig. 5A). Groups of mice were immunized with two dosages of CH-17-D/ZIKV and 17-D vaccine strain twenty-one days prior to challenge with a ZIKV African strain (Dak84). Three control groups were also used: one immunized with PBS and then challenged (named unvaccinated group), one immunized with ZIKV PF and then challenged (named ZIKV PF group) and one immunized and challenged with PBS (mock). Since 100% of the mice of the control group ZIKV PF were viremic at day 2 and 3 post-challenge, this criterion was not used to assess protection (Supplemental Fig. 3 and Supplemental Table 1). Thereby, the protection was evaluated by determining the proportion of mice with organs (brain and spleen) positive for the presence of ZIKV at day 10 post-challenge. We observed that 10% of the spleens and brains from mice immunized with the chimeric virus (both groups) were positive (Table 2). In contrast, respectively 100% and 87.5% of the spleens and brains from mice immunized with the 17-D vaccine strain (both groups) were positive *(p* value = 0.0004 for spleens and 0.0029 for brains; Fisher exact test). As expected 100% and 0% of the organs respectively from mice of the unvaccinated group and from the ZIKV PF group were positive. Viral RNA yields found in the organs were quite variable in all positive samples (Fig. 5B). These results demonstrated that immunization with the chimeric virus significantly protected mice against the systemic and brain infection induced by a heterologous ZIKV strain.

**Figure 5.**
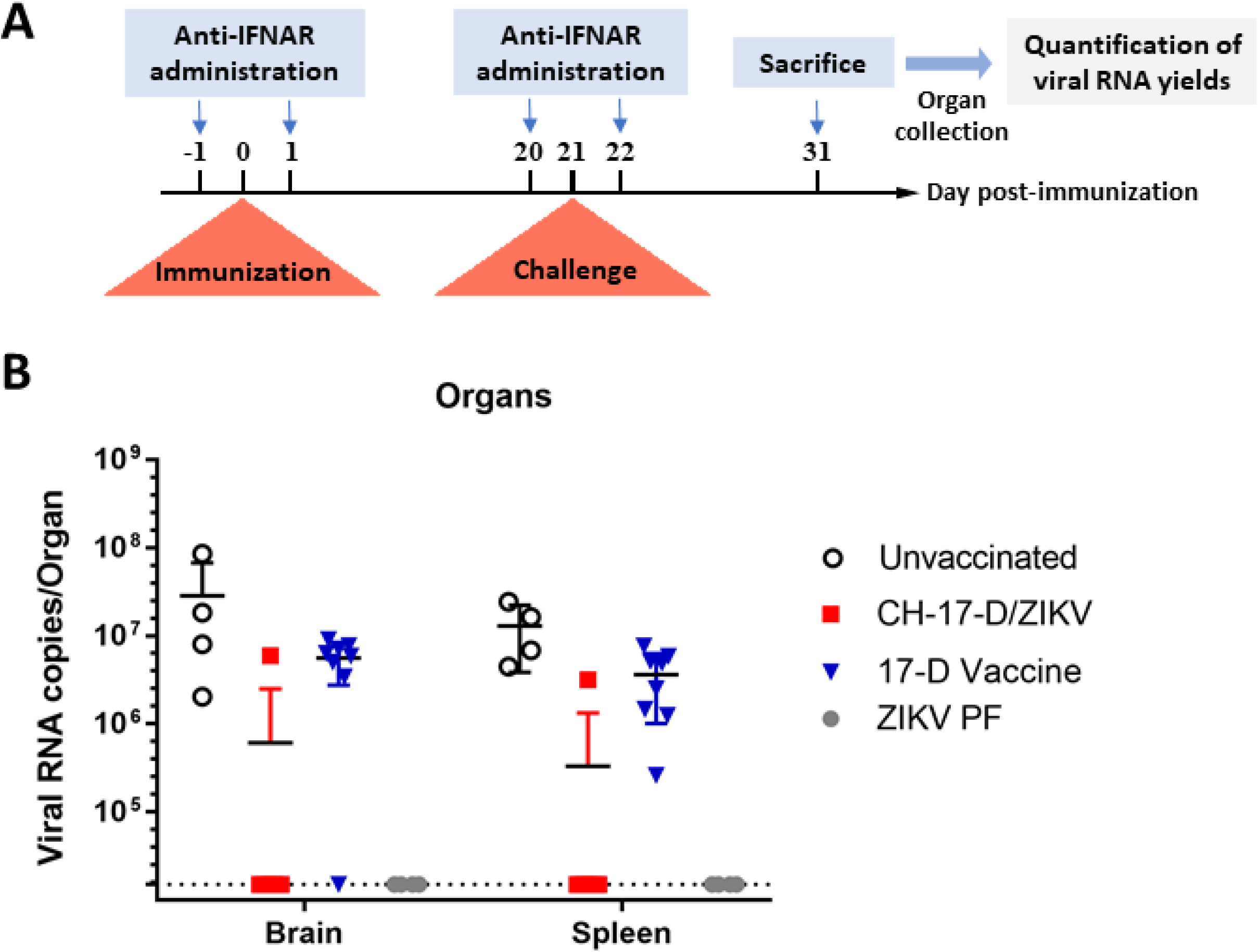
Amounts of viral RNA detected in brain and spleen samples collected during challenge experiments. **Panel A**: Experimental timeline. **Panel B**: Amounts of viral RNA in brain and spleen samples collected during challenge experiments (cf. Table 2) measured using a real time quantitative RT-PCR assay. Mean values ±SD are represented respectively by black lines and error-bars. Results from both doses of viruses are pooled.

**Table 2:** Protection of transitory immunocompromised mice challenged with a heterologous strain of ZIKV. Groups of mice were immunized with two doses (10e4 and 10e5 TCID50) of CH-17-D/ZIKV, the 17-D vaccine strain or PBS (unvaccinated). Twenty-one days later, mice were challenged with 10e6 TCID50 of a ZIKV African strain. The proportion of mice with positive spleen/brain at day 10 post-challenge was expressed as percentage. Results from both doses of viruses are pooled (results for each group are in supplemental Table 2). Detection of viral RNA was assessed using a real time RT-PCR assay. Amounts of viral RNA detected in samples are represented in figure 5.

| Viral strain | Spleens | Brains |
| --- | --- | --- |
| CH-17D/ZIKV (both doses) | 10% (1/10) | 10% (1/10) |
| 17-D vaccine strain (both doses) | 100 % (8/8) | 87,5% (7/8) |
| Unvaccinated | 100% (4/4) | 100% (4/4) |
| ZIKV PF | 0% (0/4) | 0% (0/4) |

## Discussion

We present here the initial development of a chimeric ZIKV live-attenuated vaccine candidate based on a yellow fever attenuated 17-D genetic backbone. Using the ISA reverse genetics method, we were able to rapidly test several associations of subgenomic amplicons starting from a pre-existing 17-D vaccine strain reverse genetics system. This method was recently applied to Asian and African strains of ZIKV^27^.

Three different designs were tested to incorporate the prM/E of ZIKV in the 17-D vaccine genetic backbone. Our results highlighted the necessity of modifying the cleavage site between the pre-peptide and prM protein for the construction of chimeric viruses as already described during development of chimeric ZIKV/DENV and DENV/ZIKV strains^28^.

Nevertheless, we also demonstrated that chimeric viruses needed to acquire adaptive mutations to properly replicate in mammalian cells. Indeed, we observed low-percentage of virus recovery during cell transfection experiments and both replicative viruses rescued shared five substitutions of which two were non-synonymous and located in domain II of the E protein. Interestingly, mutations located in this particular domain of the E protein were previously described *in cellulo* with 17-D vaccine strain based chimeric flaviviruses (DENV type 1/2 and Japanese encephalitis virus)^29,30^. These findings suggest that emergence of compensatory mutations in the E protein is probably necessary to restore the replicative fitness of the virus following exchange of two of its structural proteins.

By comparing the growth properties of our chimeric virus with its two parental strains in different mammalian cells, we found that this new synthetic virus had its own biological properties probably in relation to the nature of this new association of viral proteins. In fact, we observed than this strain is fitter than parental strains in Vero-E6 cells and close to 17-D vaccine strain in HEK-293 cells.

Genetic stability is a major concern when designing future live-attenuated vaccine candidates. Using Vero cells, widely used for vaccine production^22^ and the adapted chimeric virus, we performed serial passages to assess this essential criterion. We demonstrated that once initial adaptation was achieved, the chimeric virus remained genetically stable.

We used transitory immunocompromised mice as an animal model system to characterize *in vivo* the chimeric virus. We demonstrated that mice infected with this virus produced levels of neutralizing antibodies close to those observed following infection by ZIKV. Our results also showed that immunization using the chimeric strain significantly protected mice against brain and spleen invasion induced by a heterologous strain of ZIKV. Altogether, these results provide evidence that this chimeric strain had all the prerequisites to be tested in a more relevant animal model such as the microcephalic sensitive mouse model^31^.

The strategy used in the present study to develop a live ZIKV vaccine candidate has several advantages: the 17-D vaccine strain has long history of use in hundreds of millions of persons^32^ and is considered as the safest live-attenuated vaccine^33^. Moreover, in comparison with targeted attenuation strategies such as local modification of genomic region, our approach eliminates the risk of phenotype reversion by potential homologous recombination. Finally, although potential occurrence of antibody dependent enhancement phenomenon consecutive has to be considered with chimeric vaccine, there is currently no epidemiological data supporting this hypothesis in area where several flaviruses co-circulate^34^.

In conclusion, our data provide sound basis for the future development of this vaccine candidate. Furthermore, the approach used in this study to rescue the chimeric virus showed that significant advances in the development of reverse genetics methods now offer the possibility of drastically reducing the timeframe between the emergence of a novel viral pathogen and the availability of a live-attenuated vaccine candidate.

## Materials and Methods

### Cell lines

All cells were grown at 37 °C in 5% CO2 with 1% Penicillin/Streptomycin (PS; 5000U.mL-1 and 5000μg.mL-1; Life Technologies) and supplemented with 1% non-essential amino acids (Life Technologies) in media as specified below.

Baby hamster kidney (BHK-21; ATCC number CCL-10), human hepatocellular carcinoma (HUH7.5; ^35^) and human embryonic kidney (HEK-293; ATCC number CCL-1573) cells were grown in Dulbecco’s Modified Eagle’s Medium High glucose (4500 mg/l) (Life Technologies) with 7.5% heat-inactivated fetal bovine serum (FBS; Life Technologies). Vero (ATCC number CCL-81) and Vero-E6 (ATCC number CRL-1586) cells were grown in minimal essential medium (Life Technologies) with 7.5% FBS.

### Viruses

ZIKV Asian lineage strains PF (H/PF/2013, GenBank accession number: KJ776791) and Mart2015 (MRS_OPY_Martinique_PaRi_2015, GenBank accession number: KU647676), ZIKV African lineage strain Dak84 (A.taylori-tc/SEN/1984/41662-DAK, GenBank accession number: KU955592), YFV 17-D strain (produced by reverse genetics as described below; GenBank accession number: EU074025) and YFV strain BOL 88/1999 (isolated in 2009 from human serum and kindly provided by the National Center of Tropical Diseases (CENETROP), Santa-Cruz, Bolivia, GenBank accession number: KF907504) were used in this study. All these viral strains are available for the scientific community via the European Virus Archive goes Global (EVAg) project, a non-profit organization (https://www.european-virus-archive.com).

For each viral strain, we prepared a stock solution of clarified cell culture medium which was then used for all analyses. Briefly, a 25 cm2 culture flask of confluent Vero-E6 cells containing 667μL medium with 2.5% FBS (Life Technologies) was inoculated with 333μL of clarified infectious medium, incubated for 6 hours, washed once with Hank’s Balanced Salt Solution (HBSS, Life Technologies) and incubated for 3 days with 7mL of fresh medium. Cell supernatant medium was harvested, clarified by centrifugation, supplemented with HEPES buffer (Final concentration of 25mM; Sigma), aliquoted and stored at -80°C.

All experiments using replicating viruses were performed in BSL3 facilities.

### ISA procedure

Chimeric viruses and the YFV 17-D vaccine strain were rescued using the ISA (Infectious Subgenomic Amplicons) reverse genetics method as previously described ^20,27,36^.

#### Preparation of subgenomic DNA fragments

The complete viral genome was amplified by PCR as three overlapping DNA fragments. The first and last fragments were flanked respectively by the 5’ and 3’ termini and included the human cytomegalovirus promoter (pCMV) and the hepatitis delta ribozyme followed by the simian virus 40 polyadenylation signal (HDR/SV40pA). We started by using a reverse genetics system designed for the YFV 17-D strain (available on supplemental material). Because the first DNA fragment contained all regions encoding structural genes, only this fragment was modified to design chimeric viruses (the primers are listed in supplemental table 3).

DNA fragments were generated by PCR using *de novo* synthesized genes (Genscript) as templates. The sequence of the primers used is listed in supplemental table 2. PCR mixes were prepared using the Platinum PCR SuperMix High Fidelity kit (Life Technologies) following the manufacturer’s instructions. Amplifications were performed using ABI 2720 thermal cycler (Applied Biosytems) with the following conditions: 94°C for 2 min followed by 40 cycles of 94°C for 15 s, 60°C for 30 s, 68°C for 5 min plus 10 min final elongation at 68°C. PCR product size and quality were controlled by running gel electrophoresis and DNA fragments were purified using the High pure PCR product purification kit (Roche).

#### Cell transfection

Mixtures of BHK-21 and HEK-293 cells were seeded into PureCoat amine 6-well cell culture plates (Corning) one-day prior to transfection. Cells were transfected with 2μg of an equimolar mix of the three DNA fragments using lipofectamine 3000 (Life Technologies) and following the manufacturer’s instructions. Each transfection was performed in 5 replicates. After 24h of incubation, the cell supernatant media were removed and replaced by fresh cell culture medium. Six days post-transfection, cell supernatant media were passaged four times using 6-well cell culture plates of confluent Vero-E6 cells: cells were inoculated with 100μL of diluted (1/3) cell supernatant media, incubated 2h, washed with HBSS and incubated six days with 3mL of medium. Remaining cell supernatant media were stored at -80°C and called passage #1, #2, #3 and #4. To ensure the complete removing of DNA used during the transfection, the passage #4 was used to assess virus replication: 100μL of cell supernatant medium was collected to detect viral RNA using a qRT-PCR assay (see below). Passage #3 was used to produce virus stock solutions of YFV 17-D and chimeric viruses.

### RNA extraction and real-time quantitative PCR assays

RNA extraction was performed using the Qiacube HT and the Cador pathogen extraction kit (both from Qiagen) following the manufacturer’s instructions. Briefly, 100μl of cell supernatant medium was collected into an Sblock containing recommended quantities of VXL, proteinase K and RNA carrier. A DNAse digestion step (QIAGEN) was performed to remove the DNA used during cell transfection. The quantity of viral RNA was quantified by real-time quantitative RT-PCR (qRT-PCR; EXPRESS One-Step Superscript™ qRT-PCR Kit, universal, Life Technologies). The sequence of the primers used to detect ZIKVs, YFV 17-D and chimeric viruses is listed in supplemental table 4. 3.5 μL of RNA was used for each reaction (final volume of 10μL). Amplifications were performed using the QuantStudio 12K Flex Real-Time PCR System (Applied Biosytems) with the following conditions: 10 min at 50°C, 2 min at 95°C, and 40 amplification cycles (95°C for 3 sec followed by 30sec at 60°C). Amounts of viral RNA were calculated from standard curves (quantified T7-generated synthetic RNA standards were used).

### Complete Genome sequencing

Complete and partial genome sequencing of chimeric virus were performed as previously described ^37^ Viral RNA extraction was performed as described above. A set of specific primer pairs (supplemental table 5) was used to generate amplicons by RT-PCR using the Superscript III One-Step RT-PCR Platinum TaqHifi kit (Life Technologies). For each passage sequenced, purified PCR products were pooled and analyzed using the Ion PGM Sequencer (Life technologies) according to the manufacturer’s instructions. Resulting reads were analyzed using the CLC Genomics Workbench 6 software (CLC Bio). They were trimmed using quality score, by removing the primer sequences at their termini and by finally systematically removing 20 nt at the 5’ and 3’ termini. Remaining reads with a length greater than 99 were mapped using the designed sequence of the chimeric virus as reference to obtain a consensus sequence. Mutation frequency for each position was calculated as the number of mutated reads divided by the total number of reads at that site.

### Tissue culture infectious dose 50 (TCID50) assay

96-well cell culture plate of confluent Vero-E6 containing 100μL/well of media was inoculated (50μL/well) with 10-fold serial dilutions of centrifugation clarified cell culture supernatant medium. Each dilution was repeated 6 times. The plate was incubated for 7 days and read for absence or presence of CPE in each well. TCID50 titers were then calculated using the Reed-Muench method ^38^.

### Cell Viability assay

Confluent cells were inoculated at an MOI (multiplicity of infection) of 0.01 in a 96-well cell culture plate in triplicate for each measurement day. Every day for a period of 5 days we performed the cell titer blue viability assay (Promega) following the manufacturer’s instructions.

### Virus growth kinetics

Confluent cells were inoculated at an MOI of 0.01 in a 6-well cell culture plate in triplicate. Every day for a period of 7 days, 100μL of cell supernatant medium was collected to measure amounts of viral RNA by qRT-PCR (see above) and 200μl for TCID50.

### Indirect immunofluorescence assay

Confluent Vero-E6 cells were inoculated at an MOI of 0.01 in an 8-well cell culture in Lab-Tek II Chamber Slide System in duplicate. At 2 and 5 days post-infection, cells were washed twice with HBSS and fixed with 4% paraformaldehyde for 2 hours. Detection of viral antigens was performed as previously described^20,39^ using a specific ZIKV immune serum as the primary antibody (dilution: 1/50) collected from a Syrian Hamster immunized with the ZIKV strain Mart2015 (see below). This serum was shown to neutralize more than 90 % of ZIKV PF replication until 1/3000 dilution (data not shown). The secondary antibody used was a Goat anti-hamster Alexa 488 antibody (Invitrogen) at 1/500 dilution. Slides were observed using a Eurostar II fluorescence microscope with the Europicture software (Euroimmune).

### Viral RNA Yield Reduction Neutralization Test (YRNT)

Vero-E6 cells were seeded into a 96-well cell culture plate one day prior to infection (5×10^4^ cells in 100μL of medium containing 2.5% FBS per well). The next day, two fold serial dilutions of sera (from 1/20 to 2560; diluted with medium containing 2.5% FBS) were mixed (50:50; v/v) with appropriate amounts of viral stock (diluted in medium containing 2.5% FBS), incubated 1h 30 min. at 37°C/5%CO2 and then added to cells (50μL/well). The amount of virus added had been calibrated to ensure that virus production in cell supernatant medium did not reach a plateau at the readout time^40^. Cells were incubated for 3 days and 100μL of cell supernatant medium was harvested to perform nucleic acid extraction and to quantify amounts of viral RNA using a real-time qRT-PCR assay (see above). Each serum dilution was tested in triplicate and duplicate for the control group. For each experiment, a virus replication control (VC) was performed in quadruplicate to asses viral replication. For each serum dilution, viral RNA yield reduction (% of viral inhibition) was calculated using as reference the mean amount of viral RNA obtained with VC. 50% and 90% viral inhibition cut-offs were used to estimate ‘viral RNA Yield Reduction Neutralization 50% and 90% (YRNT50 ; YRNT90) titers using the method of Reed and Muench^38^.

### *In vivo* experiments

#### Ethics statement

Animal protocols had been approved by the local ethics committee (Comité d’éthique en expérimentation animale de Marseille - C2EA -14; protocol number #9327). All *in vivo* experiments were performed in accordance with the European legislation covering the use of animals for scientific purposes (Directive 210/63/EU) and French national guidelines.

#### Animal handling

Animals were maintained in ISOcage P Bioexclusion System (Techniplast) with unlimited access to food and water and 12h-light/12h-dark cycle. Animals were individually monitored every day to detect appearance of any clinical sign of illness/suffering. Virus/Antibody inoculation, blood collection as well as euthanasia (cervical dislocation) were performed under general anesthesia (isofluorane).

#### Golden hamster immunization

One 4-weeks-old female Syrian Hamster (Janvier) was intra-peritoneally immunized with 100μL containing 10^5^ TCID50 of ZIKV strain Mart2015. 21 days later the Hamster was reinjected with the same dose. The hamster did not show any sign of illness or weight loss. 15 days later, the hamster was euthanized and a blood sample (intracardiac puncture) was collected. After centrifugation, serum was stored at -80°C.

#### Administration of anti-IFNAR antibody

All the mice used were immunocompromised following a two-step inoculation of anti-IFNAR antibody (clone MAR1-5A3; Interchim; intraperitoneal injection ; 120μL)^23,25^: 1mg one day prior and 1 mg one day after each infection/immunization (i.e. the mice challenged were immunocompromised twice with this two-step procedure).

#### Mice immunization

Six groups of four 3-weeks-old female C57/bl6 mice (Charles River) were intraperitoneally inoculated with 100μL of virus: two groups were immunized with the YFV 17-D strain (two dosages: 10^4^ and 10^5^ TCID50), two groups were immunized with the ZIKV PF strain (two dosages: 10^5^ and 10^6^ TCID50) and two groups were immunized with the CH-17-D/ZIKV strain (two dosages: 10^4^ and 10^5^ TCID50). A control group of four mice was used as negative control group (non-immunized mice).

Blood collection (10μL) from the tail vein was performed at days 2 and 3 post-immunization to detect infectious virus by cell culture isolation. Immediately after its collection, all the blood was inoculated into a 12-well cell culture plate containing confluent Vero-E6 cells and 150μL of medium/well. After an incubation of 2 hours, 100μL of the inoculum was harvested. The cells were washed (HBSS), 1.5mL/well of fresh medium was added to the cells which were incubated for 6 days. Finally, 100μL of cell supernatant medium was harvested to perform nucleic acid extraction and to quantify amounts of viral RNA using a real-time qRT-PCR assay as described above.

At day 21 post-infection, all animals were euthanized. Blood samples were then collected (intracardiac puncture) from euthanized animals. After blood centrifugation, sera were stored at -80°C before being used to perform the neutralization tests.

#### Challenge experiments

Five groups of four 3-weeks-old female C57/bl6 mice (Charles River) were intraperitoneally inoculated with 100μL of virus: two groups were immunized with the YFV 17-D strain (two dosages: 10^4^ and 10^5^ TCID50), two groups were immunized with the CH-17-D/ZIKV strain (two dosages: 10^4^ and 10^5^ TCID50), and one group was immunized mice with the ZIKV PF strain (10^5^ TCID50). Two control groups of 4 mice were used to perform (i) mock control group (non-immunized/non-challenged mice); and (ii) negative control group (non-immunized mice; challenged.

All animals (except mock control group) were then challenged with 10^6^ TCID50 of ZIKV Dak84. Blood collection (10μL) from the tail vein was performed at days 2 and 3 post-challenge to assess viremia by qRT-PCR. At day 10 post-challenge, all the animals were euthanized. Organs (spleen and brain) were then collected in 1 mL of HBSS supplemented with 10% of FBS and crushed 10 minutes at 30 cycles per second with one bead of tungsten using the Tissue Lyser machine (Retsch MM400). After centrifugation at 5000g during 10 min, the supernatant medium was collected centrifuged again at 10,000g for 10 min. 50 μL of the supernatant medium was used to perform nucleic acid extraction and to quantify amounts of viral RNA using a real-time qRT-PCR assay (see above).

### Statistical analysis

All data obtained were analyzed using Graph pad prism 7 software (Graph pad software). All graphical representations and statistical analyses were also performed on Graph pad prism 7 software.

## Acknowledgements

We thank Ernest A Gould from the UMR UVE (Marseille, France) for his valuable review of the manuscript. We also thank Geraldine Piorkowski and Karine Barthelemy from the UMR UVE (Marseille, France) for all the viral sequencing. This work was supported by the French “Agence Nationale de la Recherche” (grant agreement no. ANR-14-CE14-0001) and the European Virus Archive goes global project (EVAg; European Union – Horizon 2020 programme under grant agreement no. 653316; http://www.european-virus-archive.com/).

## Contributions

FT, XDL and AN conceived the experiments. XDL allowed the funding of this study. FT, MG, FA and RK performed the experiments. FT, MG, FA, RK and AN analyzed the results. FT, MG and AN wrote the paper. FA, RK and XDL reviewed and edited the paper.

